# Aging dampens the intestinal innate immune response during *Clostridioides difficile* infection and is associated with altered intestinal eosinophil mobilization

**DOI:** 10.1101/2020.01.02.893461

**Authors:** Lisa Abernathy-Close, Michael G. Dieterle, Kimberly C. Vendrov, Ingrid L. Bergin, Vincent B. Young

## Abstract

*Clostridioides* (formerly *Clostridium*) *difficile* is the most common cause of hospital-acquired infection, and advanced age is a risk factor for *C. difficile* infection. Disruption of the intestinal microbiota and immune responses contribute to host susceptibility and severity of *C. difficile* infection. However, the impact of aging on the cellular immune response associated with *C. difficile* infection in the setting of advanced age remains to be well described. This study explores the effect of age on cellular immune responses in *C. difficile* infection as well as disease severity. Young adult mice (2-3 months old) and aged mice (22-28 months old) were rendered susceptible to *C. difficile* infection with cefoperazone and then infected with *C. difficile* strains of varying disease-causing potential. Aged mice infected with *C. difficile* develop more severe clinical disease, compared to young mice. Tissue-specific CD45+ immune cell responses occurred at the time of peak disease severity in the cecum and colon of all mice infected with a high-virulence strain of *C. difficile*; however, significant deficits in intestinal neutrophils and eosinophils were detected in aged mice. Interestingly, while *C. difficile* infection in young mice was associated with a robust increase in cecal and colonic eosinophils, there was a complete lack of an intestinal eosinophil response in aged counterparts accompanied by a simultaneous increase in blood eosinophils with severe disease. These findings demonstrate that age-related alterations in immune responses are associated with significantly worse *C. difficile* infection and support a key role for intestinal eosinophils in mitigating *C. difficile*-mediated disease severity.

## INTRODUCTION

In the last two decades, the frequency of *C. difficile* infection (CDI) among hospitalized patients has steadily increased, particularly among those 65 years of age and older (1). Several studies have demonstrated that as an individual’s age increases, so does their risk of *C. difficile* infection and the severity of CDI-associated disease (2, 3). While the connection between advanced age and severe CDI disease outcomes has been well established, the contribution of the aging host’s immune responses during acute CDI disease development and pathogenicity of *C. difficile* strain remains to be clarified.

Recently, peripheral eosinophil counts were found to be predictive of CDI disease severity and mortality in patients (4), and eosinophils were shown to potentially be protective in mouse models of CDI (5, 6). Eosinophils are innate immune cells that predominantly reside in close proximity to microbes that colonize mucosal surfaces under non-inflammatory homeostasis (7). The biological function of eosinophils in health and disease are most well studied and described in the protection against helminth infections (8) and in the pathogenesis of allergy (9). There is now growing evidence supporting a previously under-appreciated role for eosinophils as important mediators of intestinal immune responses (10), and the expression of a broad range of pattern-recognition receptors in eosinophils suggest a potential role in bacterial infection (11). Recent efforts by multiple research groups have indicated a role for eosinophils in CDI disease (4–6). However, the specific role for eosinophils in CDI disease severity has yet to be completely elucidated, and few studies characterize the innate immune responses to *C. difficile* strains with a range of virulence potential in animals of advanced age.

CDI disease severity depends on host factors and virulence of the *C. difficile* strain *(12)*. Aging is known to cause immune dysfunction and negatively impacts patients in the setting of infectious diseases in the intestine (13). While immunosenescence likely plays a role in modulating CDI outcomes (14, 15), dysregulation of particular immune cell subsets may differentially contribute to CDI disease severity. In the present study, we characterize the effect of *C. difficile* strain virulence and host age on the cellular immune response using a murine model of CDI utilizing *C. difficile* strain VPI 10463 (high-virulence) and strain 630 (low-virulence), as well as a young cohort and an aged cohort of adult mice reared in the same animal facility.

## MATERIALS AND METHODS

### Mice

Male and female specific pathogen-free (SPF) C57BL/6 wild-type adult mice that were young (2-3 months old) or aged (22-28 months old) were used in these studies. These mice were from a breeding colon at the University of Michigan that were originally derived from Jackson Laboratories over a decade ago. Euthanasia was carried out via CO_2_ inhalation at the conclusion of the experiment. Animal studies were approved by the Institutional Animal Care & Use Committee (IACUC) at the University of Michigan and animal husbandry was performed in an AAALAC-accredited facility.

### *C. difficile* strains and growth conditions

The *C. difficile* strains used in this study include reference strain VPI 10463 (ATCC 43255) and strain 630 (ATCC BAA-1382), and have been previously described in a murine model of CDI by Theriot *et al.* (12).

### Antibiotic administration and infection with *C. difficile*

Mice were rendered susceptible to *C. difficile* infection by placing mice on 0.5 mg/mL cefoperazone (MP Pharmaceuticals) in sterile distilled drinking water (Gibco) ad libitum. The antibiotic-supplemented water was provided for 10 days, followed by 2 days of drinking water without antibiotics. Animals were then inoculated by oral gavage with 10^3^-10^4^ CFUs of *C. difficile* spores suspended in 20-100 µl of distilled water (Gibco) or mock-infected with vehicle alone. Viable spores in each inoculum was enumerated by plating for colony-forming units (CFU) per mL on pre-reduced taurocholate cycloserine cefoxitin fructose agar (TCCFA). TCCFA was prepared as originally described (16) with modifications. Briefly, the agar base consisted of 40 g of Proteose Peptone No. 3 (BD Biosciences), 5 g of Na2HPO4 (Sigma-Aldrich), 1 g of KH2PO4 (Fisher), 2 g NaCl (J.T. Baker), 0.1 g MgSO 4 (Sigma), 6 g fructose (Fisher), and 20 g of agar (Life Technologies) dissolved in 1L of Milli-Q water. The prepared medium was autoclaved and supplemented with a final concentration of 250 µg/mL D-cycloserine (Sigma-Aldrich), 16 µg/mL cefoxitin (Sigma-Aldrich), and 0.1% taurocholate (Sigma). Over the course of the experiment, mice were regularly weighed and cecal contents were collected for quantitative culture.

### *C. difficile* quantification

Cecal contents were collected in a pre-weighed sterile tube from each mouse at time of euthanasia. Immediately following collection, the tubes were re-weighed to determine fecal weight and passed into an anaerobic chamber (Coy Laboratories). Each sample was then diluted 10% (w/v) with pre-reduced sterile PBS and serially diluted onto pre-reduced TCCFA plates with or without erythromycin supplementation. *C. difficile* strain 630 is erythromycin-resistant, whereas *C. difficile* strain VPI 10463 is sensitive to erythromycin. The plates were incubated anaerobically at 37°C, and colonies were enumerated after 18 to 24 hours of incubation.

### Clinical disease severity scoring

Mice were monitored for clinical signs of disease. Disease scores were averaged based on scoring of the following features for signs of disease: weight loss, activity, posture, coat, diarrhea, eyes/nose. A 4-point scale was assigned to score each feature and the sum of these scores determined the clinical disease severity score (17).

### Lamina propria cell isolation

Cecum and colon were excised, separated, and the lumen was flushed. Residual fat was removed and tissues were opened longitudinally. Tissue was placed in pre-warm RPMI medium containing 0.5M EDTA, dithiothreitol, and fetal bovine serum (FBS) and incubated at 37°C on an orbital shaker at 150 rpm for 15 min. After incubation, a steel strainer was used to separate tissue pieces from the epithelium-containing supernatant. Tissue was minced in RPMI medium containing dispase, collagenase II, DNase I, and FBS and incubated at 37°C on an orbital shaker at 150 rpm for 30 min. Digested tissue was filtered through a 100 μm cell strainer followed by a 40 μm cell strainer. The resultant single cell suspensions were counted on a hemocytometer using trypan blue exclusion test.

### Flow cytometry

Lamina propria single-cell suspensions from colon or cecum were incubated with anti-CD16/32 to reduce non-specific binding. Cells were incubated on ice for 30 mins in the dark, with a cocktail of fluorescent antibodies consisting of anti-CD45.2 PerCP-Cy5 (clone: 104), CD3 PE (clone: 145-2C11), CD11b PE-eFluor 610 (clone: M1/70), CD11c Alexa Fluor 700 (clone: N418), Ly6G PE-Cy7 (clone: 1A8), and Siglec-F Alexa Fluor 647 (clone: E50-2440). All antibodies were purchased from eBioscience, Biolegend, or BD Biosciences. Stained cells were incubated with an eFluor 450 fixable viability dye (eBioscience) and fixed with 0.5% paraformaldehyde. Cells were acquired using a BD LSRFortessa X-20 flow cytometer (BD Biosciences, San Jose, CA) and analyzed on FlowJo v10 software (Tree Star Inc., Ashland, OR).

### Eosinophil enumeration in blood

Blood was collected via cardiac puncture in microtainer tubes with K_2_ EDTA (Sarstedt, Nümbrecht, Germany) at the experimental endpoint. Blood samples were taken immediately to the Unit for Laboratory Animal Medicine In-Vivo Animal Core and processed for a complete blood count on an automated hematology analyzer (Hemavet 950, Drew Scientific, Miami Lakes, FL).

### Statistics

One-way analysis of variance (ANOVA) with Tukey’s post-hoc test was performed using R for *C. difficile* burden and cell population analyses. Clinical scores were analyzed in python using a Kruskal-Wallis one-way ANONA with Bonferroni post-hoc test. A p-value greater p<0.05 was considered statistically significant.

## RESULTS

### Aged mice infected with a high-virulence strain of C. difficile develop more severe clinical disease compared to young mice

Young mice (2-3 months old) and aged mice (22-28 months old) were rendered susceptible to *C. difficile* infection (CDI) by treatment with the antibiotic cefoparazone prior to oral inoculation with spores derived from *C. difficile* strain 630 (low virulence) or strain VPI 10463 (high virulence) (Figure 1A). There was no age or strain-associated difference in *C. difficile* colonization (Figure 1B). At the time of peak clinical disease, *C. difficile* strain VPI 10463 causes significantly more severe disease in both young and aged mice, compared to infection with *C. difficile* strain 630 (Figure 1C, p < 0.0001).

**Figure 1.**
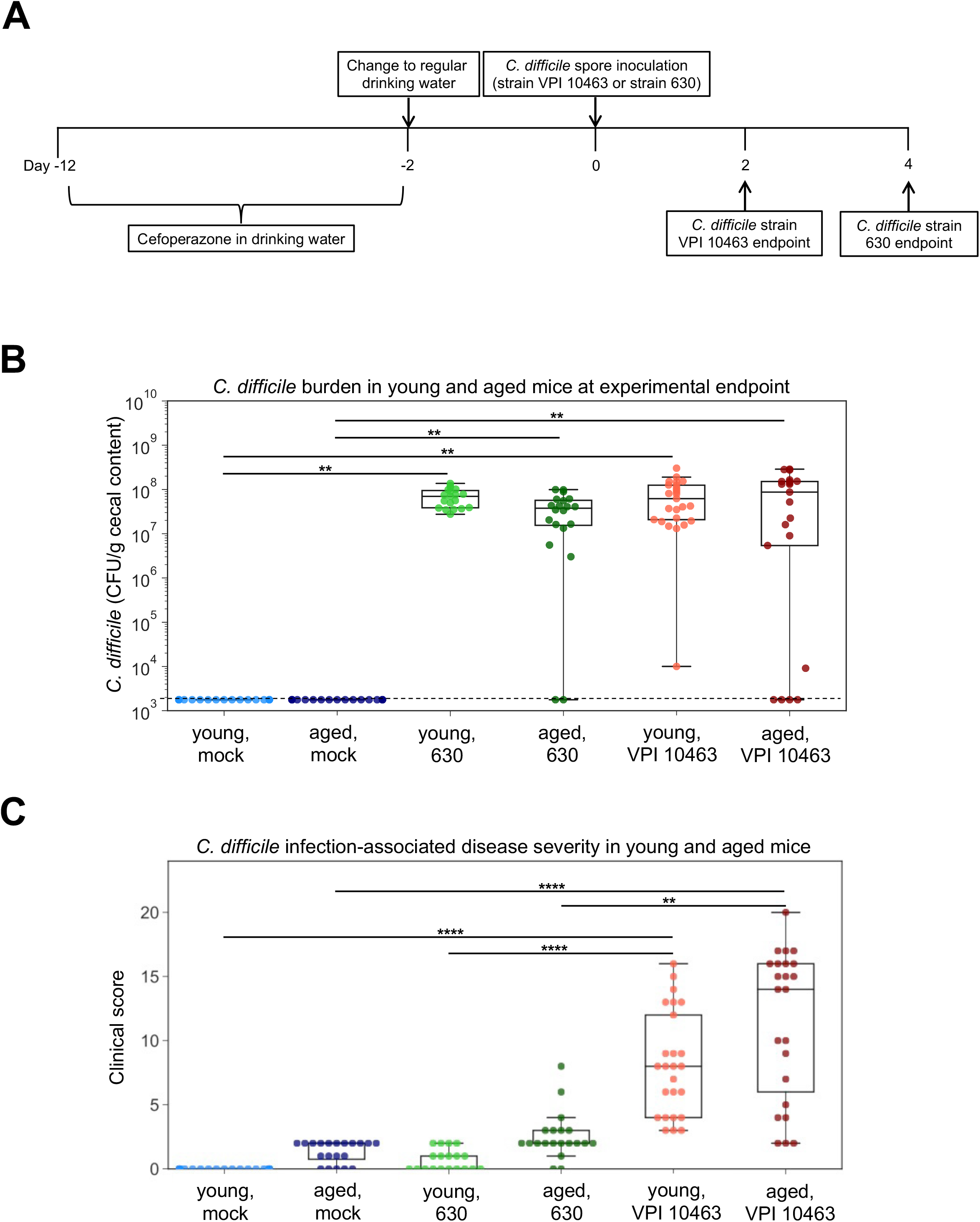
*C. difficile* infection with strain VPI 10463 or strain 630 results in increased disease severity in aged mice, compared to young mice. A) Mouse model of *C. difficile* infection in young and aged animals. B) Cecal contents were collected from young mice and aged mice at experimental endpoint (day 2 post-infection with strain VPI 10463 or day 4 post infection with strain 630) and plated anaerobically on selective agar plates to quantify *C. difficile* burden. Dotted line indicates limit of detection for *C. difficile* quantification (10^3^ CFU). C) Clinical scores of young and aged mice infected with *C. difficile* strain VPI 10463 or strain 630 at peak clinical disease severity (day 2 and 4 post-infection, respectively). One-way ANOVA with Tukey’s post-hoc test was performed for *C. difficile* burden analysis and clinical scores were analyzed using a Kruskal-Wallis one-way ANONA with Bonferroni post-hoc test, *p < 0.05; ***p <0.001.

### Infection with C. difficile strain VPI 10463, but not strain 630, results in robust cellular immune response in cecum and colon of mice, regardless of animal age

Total immune cells and myeloid cells subsets in the lamina propria of cecum and colon from young and aged mice were analyzed by flow cytometry at the time of peak disease severity (representative plots, Figure 2A and 2D). *C. difficile* strain 630 did not elicit an early cellular intestinal immune response in the cecum (Figure 2B and 2C) or colon (Figure 2E and 2F), independent of age. Due to the absence of a local intestinal cellular immune response during infection with the low-virulence *C. difficile* strain 630, we focused on further characterizing the nature of the immune cells infiltrating the distal intestinal tract during infection with the more virulent *C. difficile* strain VPI 10463.

**Figure 2.**
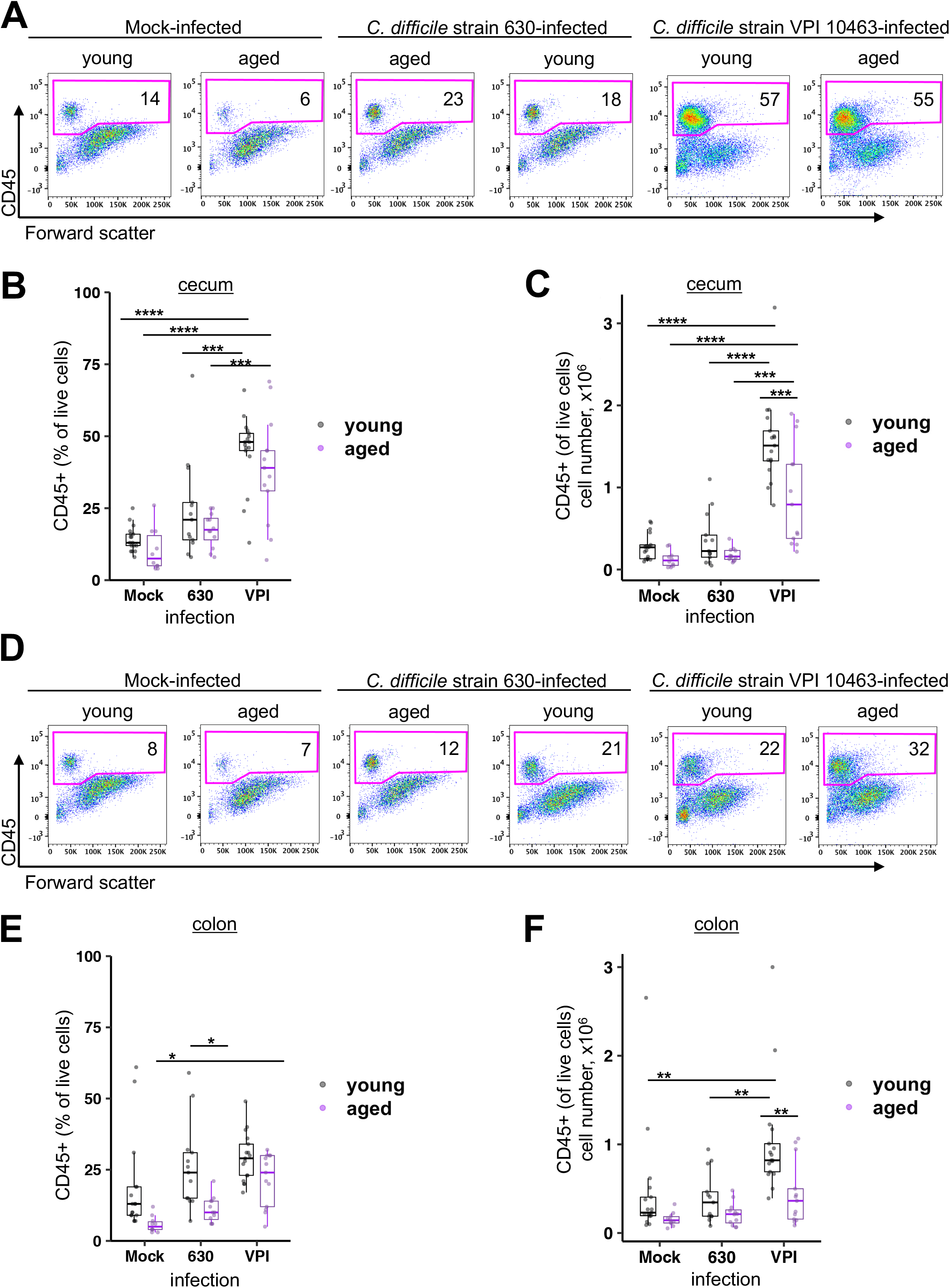
CD45+ leukocytes are preferentially increased in the cecum and colon lamina propria of young and aged mice infected with *C. difficile* strain VPI 10463, compared to infection with *C. difficile* strain 630. A) Representative flow cytometry plots indicating the percentage of CD45+ leukocytes in cecum lamina propria. B) Percentage and C) absolute number of CD45+ leukocytes in total live lamina propria cells harvested from cecum. D) Representative flow cytometry plots indicating the percentage of CD45+ leukocytes in colon lamina propria. E) Percentage and F) absolute number of CD45+ leukocytes in total live lamina propria cells harvested from colon. ANOVA and Tukey test *p<0.05; **p<0.01; ***p<0.001; ****p<0.0001.

### Aged mice mount reduced neutrophil and eosinophil responses in the distal intestine during severe C. difficile infection compared to young mice

Of the CD45+ immune cells in the cecum and colon young and aged mice infected with *C. difficile* strain VPI 10463 at the time of peak disease severity, the majority are CD11b+ cells (Figure 3A and 3B). While there is a significant increase in CD11b+ cell numbers in the cecum of young (p < 0.0001) and aged mice (p < 0.01) infected with *C. difficile* VPI 10463, this response is significantly blunted in aged mice (Figure 3C and 3D, p < 0.0001). Interestingly, we observed local differences in intestinal CD11b+ cell responses in aged mice. There was a lack of response by CD11b+ cells in the colon of aged mice infected with *C. difficile* VPI 10463, in contrast to a significant increase in colonic CD11b+ cells in young counterparts (Figure 3E and 3F). Similarly, subsets of CD11b+ myeloid cells including CD11b+Ly6G+Siglec-F-neutrophils and CD11b+Ly6G-Siglec-F+ eosinophils showed an age-dependent difference in intestinal response to *C. difficile* infection (Figure 4). While aged mice indeed mount a cecal neutrophil response during severe CDI, it is significantly dampened compared to their young counterparts (Figure 4C, p < 0.01). Furthermore, colonic neutrophil infiltration during CDI is not observed in aged mice during severe CDI (Figure 4C).

**Figure 3.**
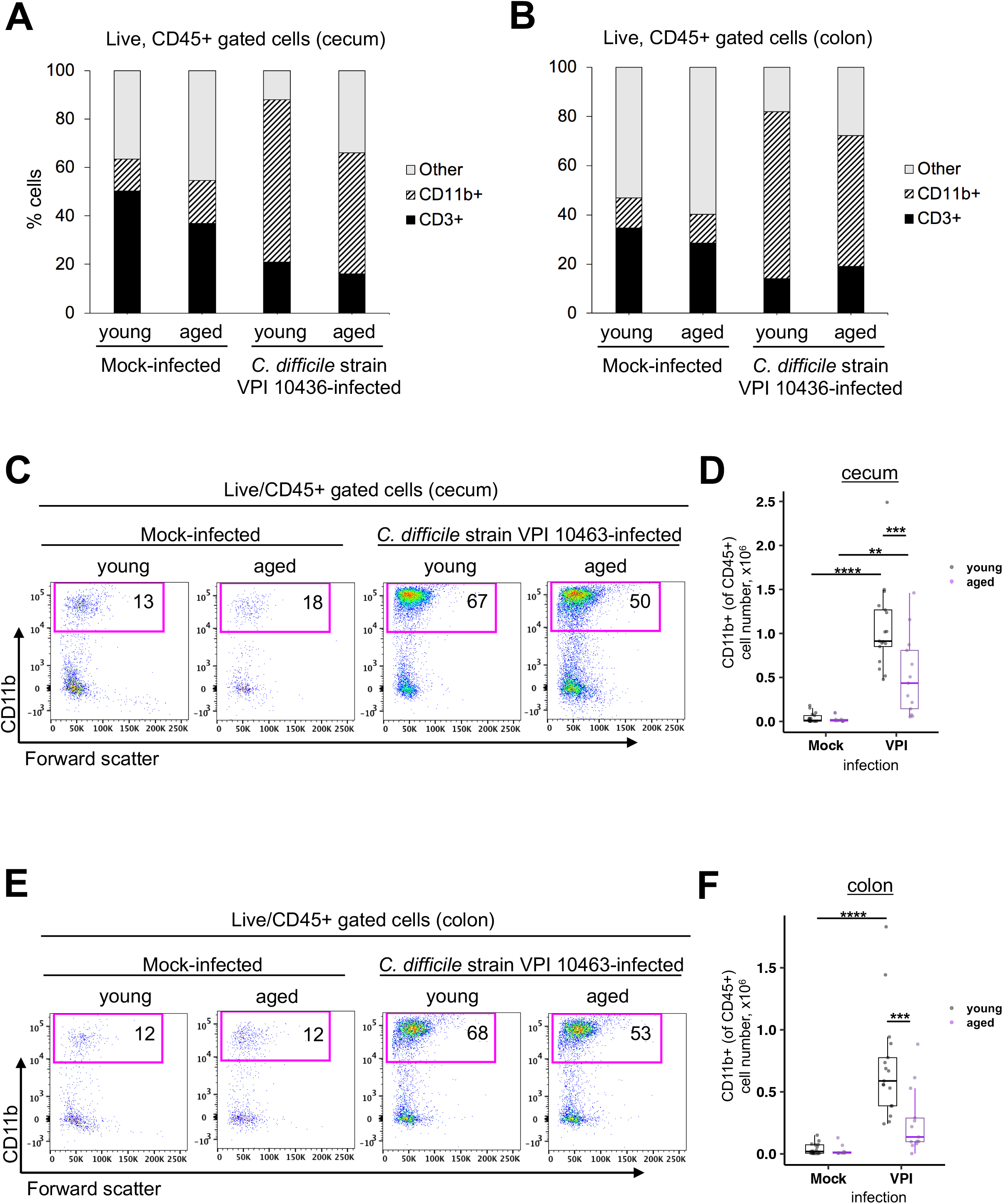
CD11b+ cells are the dominant immune cell type in the cecum and colon lamina propria of young and aged infected with *C. difficile* strain VPI 10463. Mock-infected or *C. difficile* strain VPI 10463-infected young and aged mice 2 days post-infection analyzed by flow cytometry for immune cell subsets. The ratio of CD11b+ cells, CD3+ lymphocytes, and “other” CD11b-CD3-cells of the total CD45+ immune cells in A) cecum and B) colon of mock or *C. difficile* strain VPI 10463-infected young and aged mice 2 days post-infection. C) Representative flow cytometry plots indicating the percentage of CD11b+ myeloid cells in colon lamina propria. D) Absolute number of CD11b+ cells determined by flow cytometry in cecum. E) Representative flow cytometry plots indicating the percentage of CD11b+ cells in colon lamina propria. F) Absolute number of CD11b+ cells determined by flow cytometry in colon. ANOVA and Tukey test *p<0.05; **p<0.01; ***p<0.001; ****p<0.0001.

**Figure 4.**
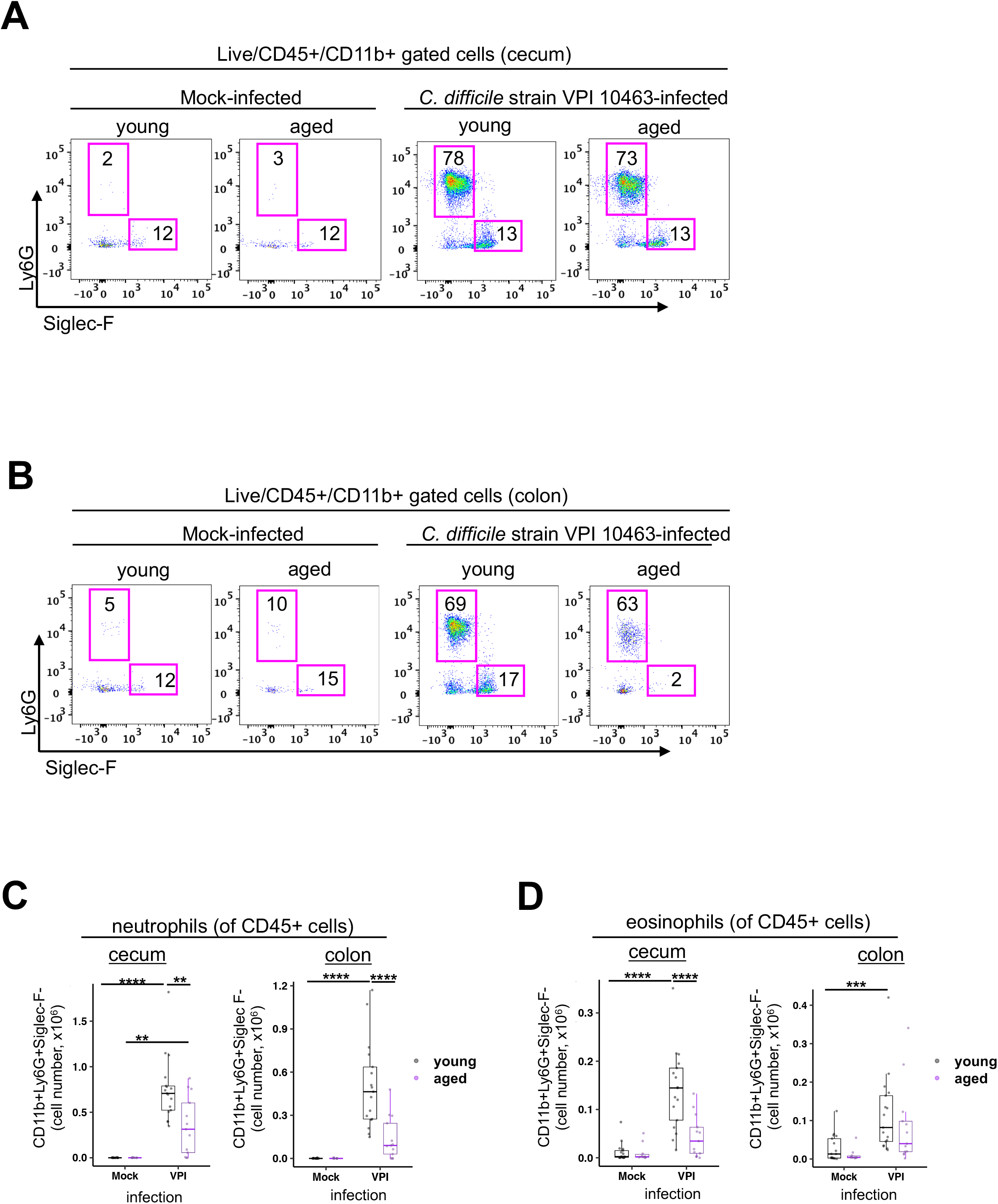
Neutrophil and eosinophil cellular infiltration in the distal intestine is significantly decreased in aged mice infected with *C. difficile* strain VPI 10463, compared to young counterparts. Mock-infected or *C. difficile* strain VPI 10463-infected young and aged mice 2 days post-infection analyzed by flow cytometry for CD11b+Ly6G+Siglec-F-neutrophils and CD11b+Ly6G-Siglec-F+ eosinophils. Representative flow cytometry plots indicating the percentage of CD11b+Ly6G+Siglec-F-neutrophils and CD11b+Ly6G-Siglec-F+ eosinophils in A) cecum and B) colon lamina propria. C) Absolute numbers of neutrophils in the cecum and colon lamina propria. D) Absolute eosinophil number in cecum and colon lamina propria. ANOVA and Tukey test *p<0.05; **p<0.01; ***p<0.001; ****p<0.0001.

### Differential intestinal and systemic eosinophil cellular response during severe C. difficile infection in aged mice

Since it has been suggested that eosinophils play a protective role in CDI using a mouse model of *C. difficile* infection (5), we hypothesized that eosinophil responses in older animals during *C. difficile* infection would differ significantly compared to their relatively young counterparts. Young mice at the time of peak CDI disease severity mounted a robust cecal and colonic eosinophil cellular response; however, intestinal eosinophil infiltration during peak CDI disease severity is absent in aged mice infected with *C. difficile* strain VPI 10463 (Figure 4D). We sought to determine if there was an age-related difference in peripheral eosinophil responses during *C. difficile* infection. We found that aged mice infected with the high-virulence *C. difficile* strain VPI 10463 respond with increased eosinophil to total leukocyte ratio (Figure 5A, p < 0.05) compared to young mice infected with the same strain of *C. difficile*. Although there is a complete lack of local eosinophil infiltration in the distal intestine of aged mice with severe CDI, there is a concomitant significant increase in the absolute number of peripheral eosinophils in the blood of aged mice infected with *C. difficile* strain VPI 10463 compared to their young counterparts (Figure 5B, p < 0.01). In contrast, young mice do not demonstrate a change in peripheral blood eosinophil levels at the peak of CDI severity, regardless of infecting *C. difficile* strain (Figure 5B). Interestingly, while severe CDI was associated with increased blood eosinophil numbers in aged mice, this was not observed with less severe CDI associated with *C. difficile* strain 630.

**Figure 5.**
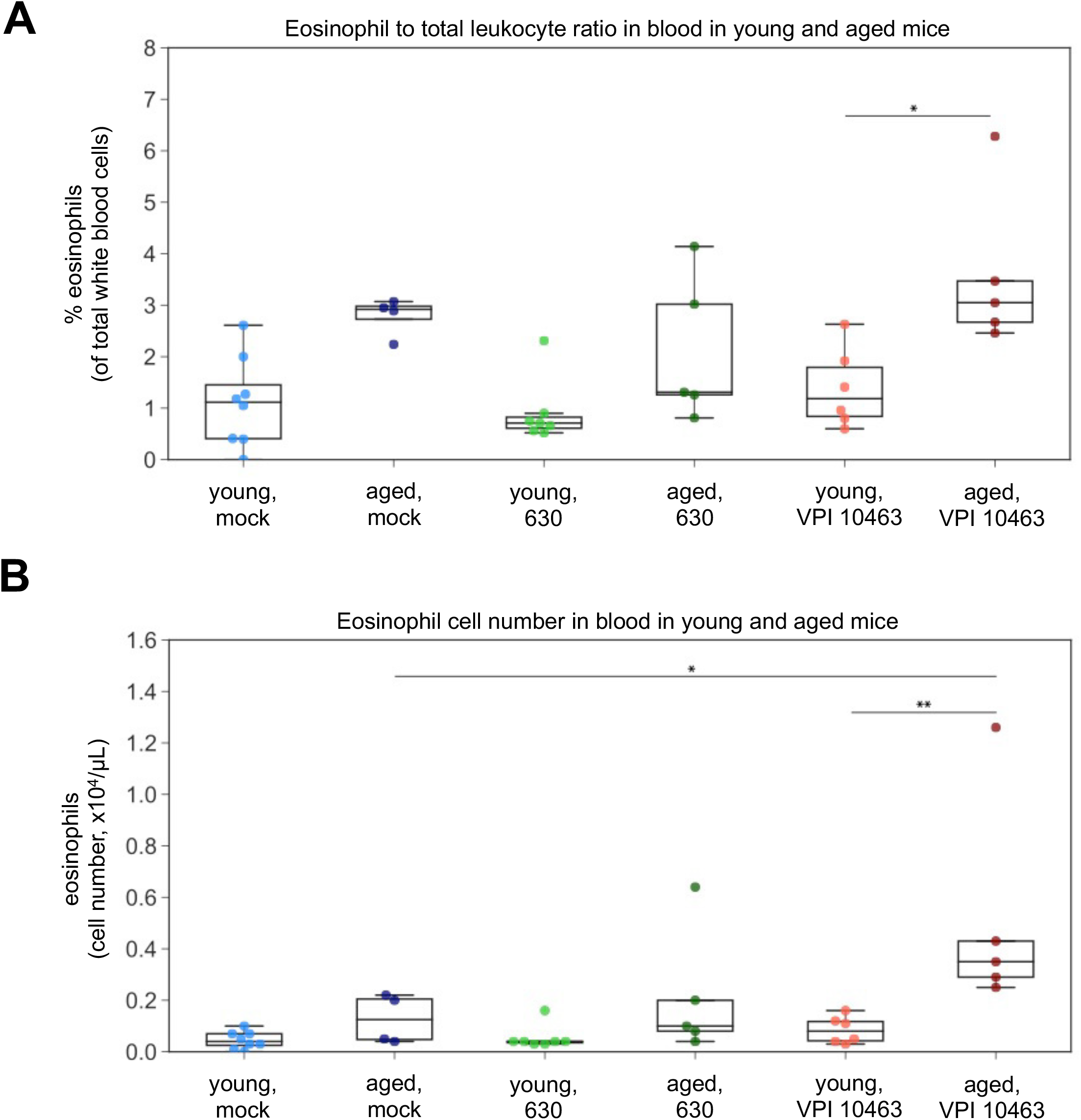
Infection with *C. difficile* strain VPI 10463 results in differential systemic eosinophil responses in aged mice, compared to young mice. A) Percentage of eosinophils of total white blood cells in young and aged mice at baseline or at peak CDI disease severity. B) Absolute numbers of eosinophils in the blood of young and aged mice at baseline or infected with *C. difficile* strain 630 or strain VPI 10463. ANOVA and Tukey test *p<0.05; **p<0.01; ***p<0.001; ****p<0.0001.

## DISCUSSION

Although advanced age is a risk factor for *C. difficile* infection (CDI) (1), the relationship between the aging immune system and *C. difficile* infection is not well known. This study demonstrates an overall decrease in intestinal innate immune responses during acute *C. difficile* infection in mice with advanced age, and identifies a specific aging-related defect in eosinophil responses during CDI. We show that aged mice (22 – 24 months old) develop more severe CDI disease and mount a significantly blunted intestinal cellular immune response compared to young mice (2 – 3 months old), with a notable absence of an intestinal eosinophil response. Interestingly, aged mice had a significant peripheral eosinophil response whereas young mice lacked an eosinophil increase detected in the blood during severe CDI. Our results also suggest that eosinophils may play differential roles, whether that be protective or pathogenic. Additionally, eosinophil counts may predict different disease outcomes during *C. difficile* infection depending on their location in intestinal tissue or circulation in the periphery. In the present study, we characterized the intestinal cellular immune response to *C. difficile* infection during peak disease severity in young and aged mice, with a focus on myeloid cell subsets mobilized during the innate immune response.

Peniche *et al.* showed that middle-aged mice (12 – 14 months old) have increased susceptibility to *C. difficile* infection and worse CDI disease, compared to young controls (18). They report that this observation was driven by impaired innate immune responses; however, eosinophils were not evaluated in this study. Our data showing a decreased intestinal neutrophil response in aged mice infected with *C. difficile* agrees with a recent study that examined the effect of age on *C. difficile infection* in a mouse model (15).

Little is known about the relationship between CDI and eosinophils. One study demonstrated that *C. difficile* toxin suppresses a host protective colonic eosinophil responses in a toll-like receptor 2 (TLR2) dependent manner (6). Recently, Buonomo *et al.* reported that an increase in intestinal eosinophils was associated with reduced host mortality during *C. difficile* infection (5). In the aforementioned study, cytokine, IgA and IgG, and *muc2* analysis were assessed in the cecum while eosinophils were enumerated in the colon. While we detected robust eosinophil infiltration in the cecum of young mice, we did not observe this in the colon of young or aged animals. Another group found that peripheral loss of eosinophils in patients with *C. difficile* infection was predictive of severe disease (4). We report that young mice had significantly increased numbers of intestinal eosinophils during *C. difficile* infection, compared to mock-infected young controls, while aged mice did not have intestinal eosinophil infiltration and significantly worse CDI disease, compared to young mice. However, eosinophils in the blood of aged mice infected with the more virulent strain of *C. difficile* were significantly increased at the time of peak CDI disease severity, compared to young counterparts. It is possible that eosinopenia at the time of symptom onset is predictive of increased CDI disease severity and mortality, but eosinophil levels may increase in the blood as CDI disease progresses over time. Additionally, eosinophil responses may be altered in older patient populations and may not be predictive of CDI outcomes. Our results also suggest tissue-specific eosinophil responses in the distal intestinal tract, so further examination of local tissue responses is warranted.

While the general blunted cellular immune response in the intestine of aged mice with CDI may be explained by immunosenescence, which describes the aging of the immune system, the observed age-related inverse of eosinophil responses to acute CDI suggests that this immune cell population plays a role in CDI pathology. Aged mice had significantly more severe clinical disease, compared to their younger counterparts. Young animals have a robust cellular intestinal immune response during acute CDI, whereas aged mice respond to *C. difficile* infection with intestinal eosinopenia and concomitant peripheral eosinophilia. These data suggest a role for eosinophils in age-associated CDI outcomes in older patient populations.

## ACKNOWLEDGMENTS

We thank Naomi Perlman and James M. George for their assistance in the laboratory, which facilitated conducting these experiments. We thank Anna Colvig and Florin Timpau of the ULAM In Vivo Animal Core for hematology assessment. This study was funded by grant U01AI12455 awarded to V.B.Y. by the National Institute of Allergy and Infectious Diseases at the National Institutes of Health. In addition, M.G.D. was supported by NIH grant T32GM007863. The funding agencies had no role in study design, data collection and interpretation, or the decision to submit the work for publication.

